# Determining the impact of putative loss-of-function variants in protein-coding genes

**DOI:** 10.1101/106468

**Authors:** Suganthi Balasubramanian, Yao Fu, Mayur Pawashe, Patrick McGillivray, Mike Jin, Jeremy Liu, Konrad J. Karczewski, Daniel G. MacArthur, Mark Gerstein

## Abstract

Variants predicted to result in the loss of function (LoF) of human genes have attracted interest because of their clinical impact and surprising prevalence in healthy individuals. Here, we present ALoFT (Annotation of Loss-of-Function Transcripts), a method to annotate and predict the disease-causing potential of LoF variants. Using data from Mendelian disease-gene discovery projects, we show that ALoFT can distinguish between LoF variants deleterious as heterozygotes and those causing disease only in the homozygous state. Investigation of variants discovered in healthy populations suggests that each individual carries at least two heterozygous premature stop alleles that could potentially lead to disease if present as homozygotes. When applied to *de novo* pLoF variants in autism-affected families, ALoFT distinguishes between deleterious variants in patients and benign variants in unaffected siblings. Finally, analysis of somatic variants in > 6,500 cancer exomes shows that pLoF variants predicted to be deleterious by ALoFT are enriched in known driver genes.

## Introduction

One of the most notable findings from personal genomics studies is that all individuals harbor loss-of-function variants in some of their genes^1^. A systematic study of LoF variants from the 1000 Genomes revealed that there are over 100 putative LoF (pLoF) variants in each individual^2–4^. Recently, a larger study aimed at elucidating rare LoF events in 2,636 Icelanders generated a catalog of 1,171 genes that contain either homozygous or compound heterozygous LoF variants with a minor allele frequency less than 2%^5^. Thus, several genes are knocked out either completely or in an isoform-specific discovery of protective LoF variants associated with beneficial traits. The potential of LoF variants in the identification of valuable drug targets has fueled an increased interest in a more thorough understanding of pLoF variants. For example, nonsense variants in *PCSK9* are associated with low LDL levels^6,7^ which prompted the active pursuit of the inhibition of *PCSK9* as a potential therapeutic for hypercholesterolemia^8^ and led to the development of two drugs which have been recently approved by the FDA. Other examples include nonsense and splice mutations in *APOC3* associated with low levels of circulating triglycerides, a nonsense mutation in *SLC30A8* resulting in about 65% reduction in risk for Type II diabetes, two splice variants in the Finnish population in *LPA* that protect against coronary heart disease, and two LoF-producing splice variants and a nonsense mutation in *HAL* associated with increased blood histidine levels and reduced risk of coronary heart disease^9–13^.

About 12% of known disease-causing mutations in the Human Gene Mutation Database (HGMD) are due to nonsense mutations^14^. pLoF variants are also prioritized in cancer studies where various filtration schemes are used to narrow down causal mutations^15,16^. Even though premature stop variants often lead to loss of function and are thus deleterious, predicting the functional impact of premature stop codons is not straightforward. Aberrant transcripts containing premature stop codons are typically removed by nonsense-mediated decay (NMD), an mRNA surveillance mechanism^17^. However, a recent large-scale expression analysis demonstrated that 68% of predicted NMD events due to premature stop variants are unsupported by RNA-Seq analyses^18^. A study aimed at understanding disease mutations using a 3D structure-based interaction network suggests that truncating mutations can give rise to functional protein products^19^. Moreover, premature stop codons in the last exon are generally not subject to NMD. Further, when a variant affects only some isoforms of a gene, it is difficult to infer its impact on gene function without the knowledge of the isoforms that are expressed in the tissue of interest and how their levels of expression affect gene function. Finally, loss-of-function of a gene might not have any impact on the fitness of the organism.

While there are several algorithms to predict the effect of missense coding variants on protein function, there is a paucity of methods that are applicable to nonsense variants^20–22^. Additionally, current prediction methods that infer the pathogenicity of variants do not take into account the zygosity of the variant^23,24^. The majority of pLoF variants in healthy population cohorts are heterozygous. It is likely that a subset of these variants will cause disease as homozygotes.

Here, we present a pipeline called ALoFT (**A**nnotation of **L**oss-**o**f-**F**unction **T**ranscripts), that provides extensive annotation of pLoF variants. Further, we developed a prediction model to classify pLoF variants into three classes: those that are benign, that lead to recessive disease (disease-causing only when homozygous) and that lead to dominant disease (disease-causing as heterozygotes). Finally, we validate the prediction model by applying ALoFT to known disease mutations in Mendelian diseases, autism and cancer.

## Results

### ALoFT pipeline

We have developed a pipeline called ALoFT to annotate pLoF variants. In this study, we include premature stop-causing (nonsense) SNPs, frameshift-causing indels and variants affecting canonical splice sites as pLoF variants, also referred to as premature truncating variants. An overview of the pipeline is shown in Supplementary Figure 1. The main features of ALoFT include (1) functional domain annotations; (2) evolutionary conservation; and (3) biological networks. For comprehensive functional annotation, we integrated several annotation resources such as PFAM and SMART functional domains^25,26^, signal peptide and transmembrane annotations, post-translational modification sites, NMD prediction^27,28^, and structure-based features such as SCOP domains and disordered residues. For evolutionary conservation, ALoFT outputs variant position-specific GERP scores, which is a measure of evolutionary conservation^29^ and dN/dS values (ratio of missense to synonymous substitution rates) for macaque and mouse that are computed from human-macaque and human-mouse orthologous alignments, respectively. In addition, we evaluate if the region removed due to the truncation of the coding sequence is evolutionarily conserved based on constrained elements^30^. ALoFT includes network features shown to be important in disease prediction algorithms: a proximity parameter that gives the number of disease genes connected to a gene in a protein-protein interaction network and the shortest path to the nearest disease gene^2,31^. The pipeline also includes features to help identify erroneous LoF calls, potential mismapping, and annotation errors, because LoF variant calls have been shown to be enriched for annotation and sequencing artifacts^2^. A detailed description of all the annotations provided by ALoFT is included in Supplementary Table 1. Documentation and GitHub link to source code can be found at aloft.gersteinlab.org.

Using the annotations output by ALoFT as predictive features (Fig. 1, Supplementary Table 2), we developed a prediction method to infer the pathogenicity of pLoF variants. To build the ALoFT classifier, we used three classes of premature stop variants as training data: benign variants, dominant disease-causing and recessive disease-causing variants (Supplementary Table 3). The benign set includes homozygous premature stop variants discovered in a cohort of 1,092 healthy people, Phase1 1000 Genomes data (1KG). Homozygous premature stop mutations from HGMD that lead to recessive disease and heterozygous premature stop variants in haplo-insufficient genes that lead to dominant disease represent the two disease classes^3,31^. In addition to loss-of-function effects, truncating mutations can also lead to gain of function. However, gain of function mutations are difficult to model systematically as the effect of variant can only be understood in the context of the biology of the gene and can vary widely for different genes and gene classes. In order to minimize errors that might arise due to inadequate modeling of gain-of-function effects and focus only on LoF, we only use predicted haploinsufficient genes as the training data for dominant model. We built the ALoFT classifier to distinguish among the three classes using a random forest algorithm^32^. For each mutation, ALoFT provides three class probability estimates, and we obtain good discrimination between each class. The average multiclass test AUC (area under the curve) with 10-fold cross-validation is 0.97. The precision for the three classes are as follows: Dominant=0.86, Recessive=0.86, Benign=0.96. The classifier is robust to the choice of training data sets and performs well with different training data sets (Supplementary Table 4, Supplementary Fig. 2). The prediction output provides three scores for each pLoF variant that correspond to the probability of the pLoF being benign, dominant or recessive disease-causing allele. In addition, ALoFT also provides the predicted pathogenicity. The pathogenic effect of pLoF variant is assigned to the class that corresponds to the maximum score. Though trained with premature stop SNVs, our method is also applicable to frameshift indels. 99.4% of HGMD disease-causing frameshift indels are predicted as pathogenic based on the maximum ALoFT score.

**Figure 1 -.**
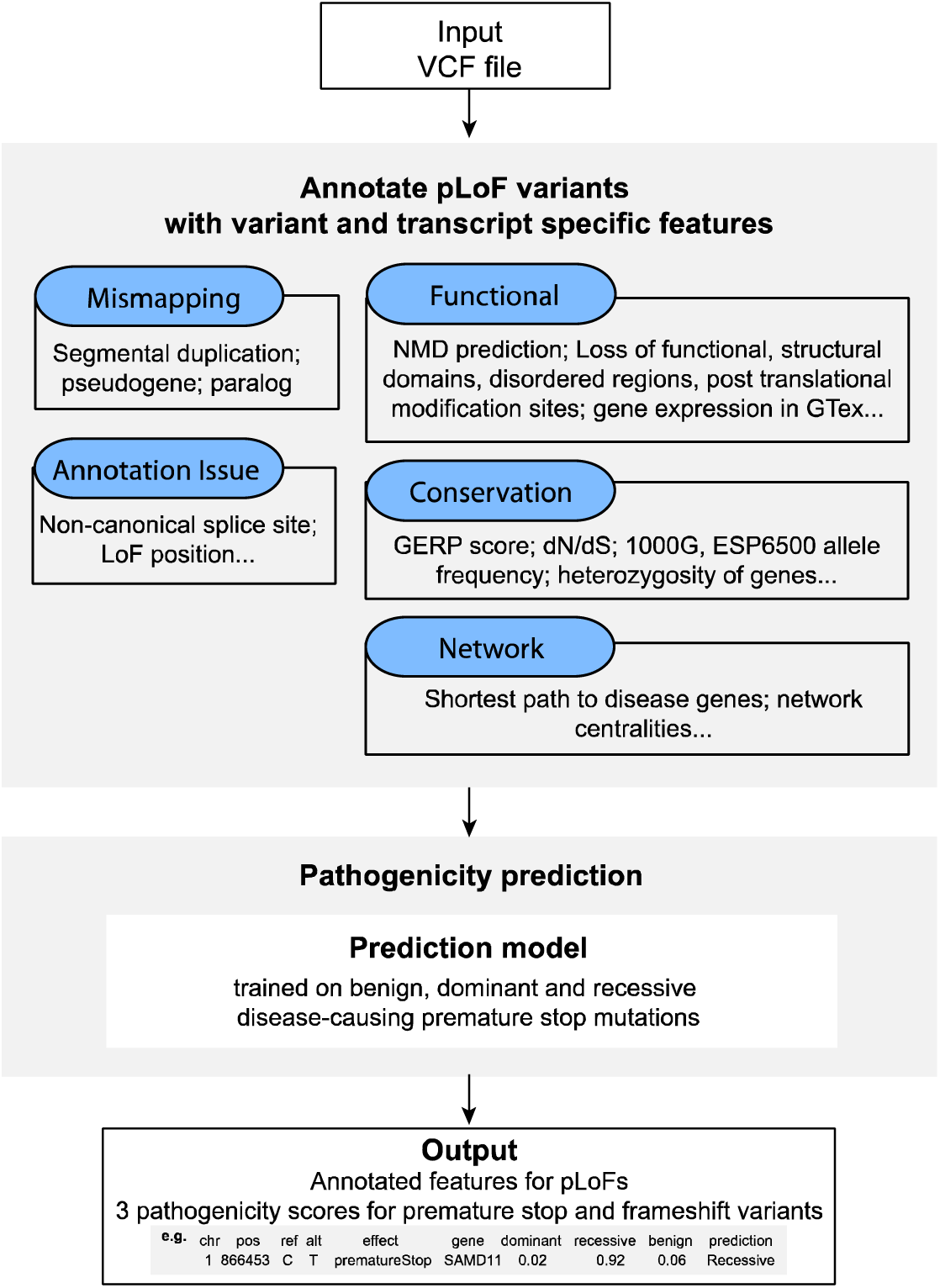
Schematic workflow. ALoFT uses a VCF file as input and annotates premature stop, frameshift-causing indel and canonical splice-site mutations with functional, conservation, network features. ALoFT also flags potential mismapping and annotation errors. Using the annotation features, ALoFT predicts the pathogenicity (as either benign, recessive or dominant disease-causing) of premature stop and frameshift mutations based on a model trained on known data. ALoFT can also take as input a 5-column tab-delimited file containing chromosome, position, variant ID, reference allele and alternate allele as its columns.

We analyzed the importance of the various features to the classification (Supplementary Fig. 3). The global allele frequency of variants in the Exome Aggregation Consortium, a dataset comprised of sequence variations obtained from an analysis of 60,706 unrelated individuals of diverse ethnicities (ExAC^33^, http://exac.broadinstitute.org), appears to be the most important feature for the classification. When we removed this feature and other features related to allele frequency (i.e. both ExAC and ESP) and retrained the random forest model, the classifier still performs well with an average multiclass test AUC of 0.93. (The precision for the three classes are as follows: Dominant=0.84, Recessive=0.80, and Benign=0.75). We also systematically evaluated the classifier using models trained on various specific sets of features (Supplementary Table 5). Overall, we find that classifier is not driven by any single feature and integrating many features improves prediction accuracy.

### Validation of classifier

We applied ALoFT to elucidate the pathogenicity of pLoF variants in various disease scenarios. Using case studies, we show that ALoFT provides robust predictions for the effect of pLoFs.

### Case study 1: Application of ALoFT to understand pLOFs in Mendelian disease

We evaluated ALoFT by predicting the effect of known disease-causing premature stop mutations from ClinVar^34^ (details in Supplementary Information) and predicted the mode of inheritance and pathogenicity of all of the truncating variants (Fig. 2a). ALoFT is clearly able to distinguish between pLoFs that possibly lead to disease in a heterozygous state versus those that do so only in a homozygous state. Our method shows that heterozygous disease-causing variants have significantly higher dominant disease-causing scores than the homozygous disease-causing variants (p-value: 1.3e-13; Wilcoxon rank-sum test). We used two other measures, GERP score, which is a measure of evolutionary conservation, and CADD score, which gives a measure of pathogenicity, to classify recessive versus dominant pLoF variants^35^. Both CADD (p-value: 0.13) and GERP (p-value: 0.49) scores are not able to discriminate between recessive and dominant disease-causing mutations (Fig. 2a). We also tested our method on a smaller dataset from the Center For Mendelian Genomics studies^36^ and was able to correctly recapitulate the pathogenic effect of pLoF variants and their inheritance pattern (Supplementary Fig. 4).

**Figure 2 -.**
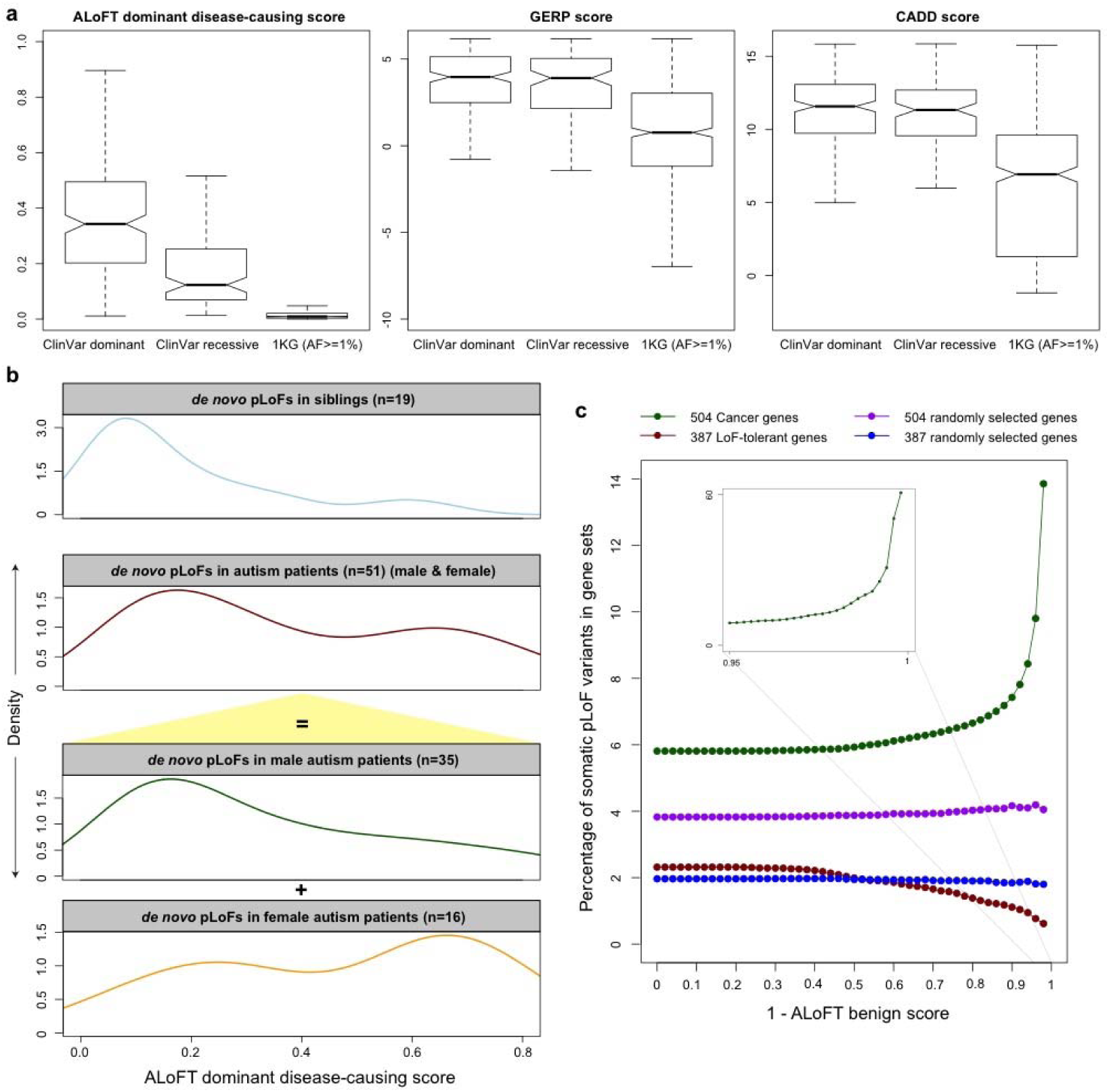
ALoFT classification of premature stop variants from Mendelian disease, autism and cancer studies. a) ALoFT dominant disease-causing score, GERP and CADD score for ClinVar and 1KG common (AF>=1%) variants. All training variants are excluded. Average tolerance scores are 0.097 and 0.115 respectively for ClinVar dominant and recessive datasets. b) The top two panels show the dominant disease-causing scores of *de novo* premature stop mutations in autism patients and siblings; mutations in patients are further separated by gender, as shown in the bottom two panels. c) The fraction of mutations occurring in various gene categories (Y-axis) as a function of predicted diseasing-causing score for cancer somatic premature stop variants (X-axis). Disease-causing score is calculated as (1- predicted benign score). We calculated the fraction of somatic premature stop mutations in 504 known cancer driver genes and 504 randomly selected genes. To ensure that the cancer driver genes and the selected random genes have similar length distributions, the 504 random genes were selected from genes with matched length. Similarly, we compared the fraction of somatic premature stop mutations in 397 LoF-tolerant genes and 397 randomly selected genes with similar length distribution. LoF-tolerant genes are genes that have at least one homozygous LoF variant in at least one individual in the 1KG cohort.

### Case study 2: Application of ALoFT to understand *de novo* pLoFs implicated in autism

*De novo* pLoF SNPs have been implicated in autism based on analysis of sporadic or simplex families (families with no prior history of autism)^37–40^. We applied our method to *de novo* pLoF mutations discovered in these studies. Our method shows that the proportion of dominant disease-causing *de novo* LoF events is significantly higher in autism patients versus siblings (Fig. 2b; p-value: 8.4e-4; Wilcoxon rank-sum test).

Autism spectrum disorder is known to be four times more prevalent in males than females suggesting a protective effect in females. Previous studies show that there is a higher mutational burden for non-synonymous mutations in females ascertained for autism spectrum disorder^41^. Therefore, we investigated the differences in the impact of *de novo* pLoF variants in male versus female autism patients. We also observe a similar pattern for pLoF mutations as has been found for missense variants – female probands have a higher proportion of predicted deleterious *de novo* pLoF variants than male probands (Fig. 2b; p-value: 0.03). A recent study based on exome sequencing of 3,871 autism cases delineated 33 risk genes at FDR < 0.1^42^. We observe that the *de novo* pLoF mutations in the autism patients in the 33 risk genes have higher dominant disease causing score than the *de novo* pLoF variants in other genes (Supplementary Fig. 5; p-value: 5e-3). Supplementary Table 6 includes the ALoFT predictions for *de novo* pLoF variants. Thus, ALoFT predictions corroborate the role of *de novo* pLoF variants in autism as shown by others using entirely different approaches.

### Case Study 3: Identification of pathogenic somatic LoF variants in cancer

We applied our prediction method to infer the effect of somatic premature stop variants (somatic pLoFs) from a compilation of 6,535 cancer exomes^43^. As shown in Fig. 2c, somatic pLoFs are enriched in known cancer driver genes compared to randomly sampled genes of matched lengths. Moreover, deleterious somatic LoFs are strongly enriched in driver genes and depleted in LoF-tolerant genes (genes that contain at least one homozygous LoF variant in the 1KG population). In the context of somatic mutations, variant zygosity, or distinguishing between ‘dominant’ and ‘recessive’ disease causing mutations, is not always relevant. Cancer cells may show aneuploidy and cellular heterogeneity. Therefore, for the evaluation of somatic mutations, we define an overall measure of deleteriousness as (1-Benign ALoFT score) on the X-axis of Fig. 2c.

We also evaluated ALoFT as a tool for distinguishing driver LoF mutations from passenger LoF mutations in tumors with high mutation burden. We observe a decrease in deleterious LoF mutations with increasing total mutational burden (Supplementary Fig. 6a). However, the ratio of deleterious LoFs to total pLoFs displayed no significant trend across groups (Supplementary Fig. 6b). The ratio of deleterious LoF mutations to total pLoF mutations is consistently high across groups (84%). This may indicate that driver LoF events tend to arise early in tumor development.

To classify genes as tumor suppressors, Vogelstein et al. proposed a “20/20” rule, whereby a gene is classified as a tumor suppressor if > 20% of the observed mutations in that gene are inactivating mutations^44^. Among the 210 genes that met 20/20 rule criteria, 87% of pLoF mutations affecting these genes were deleterious LoFs, representing 21% of total mutations. By comparison, only 6% of mutations were deleterious LoFs among 11892 genes that did not meet 20/20 criteria (p << 0.001) (Supplementary Fig. 7). A list of these genes is provided as Supplementary Table 8. In cases where genes display a high somatic pLoF rate but low somatic deleterious LoF rate, ALoFT may be used to identify potential false-positive driver genes predicted by the 20/20 rule.

### Case study 4: Distinguishing between benign and pathogenic pLoFs

Finally, we applied ALoFT to predict the effect of premature stop variants in the final exons of protein-coding genes. It is often assumed that premature stop variants in the last coding exon are likely to be benign because they could escape NMD; as a result, in many cases the effect will be the expression of a truncated protein rather than a complete loss of function. However, examples of disease-causing mutations in the last exon are also known^45^. Therefore, we applied ALoFT to see if we could distinguish between benign and disease-causing LoF variants in the last coding exon. To this end, we applied ALoFT to understand the effect of pLoF variants in ESP6500, ExAC and HGMD datasets. A higher proportion of rare variants are observed in ESP6500 and ExAC cohort due to its larger sample size and higher sequencing depth (Fig. 3a). A large number of both common and rare premature stop variants are seen at the end of the coding genes in the 1KG, ESP6500 and ExAC datasets. In contrast, fewer disease-causing HGMD variants are seen at the ends of coding genes (Fig. 3a). ALoFT predicts that both common and rare premature stop variants in the last coding exon in the 1KG, ESP6500 and ExAC cohort are likely to be benign, whereas HGMD mutations in the last coding exon tend to be disease-causing (Fig. 3b). Thus, ALoFT is able to differentiate between rare but benign premature stop variants seen in healthy individuals and the rare disease-causing HGMD alleles.

**Figure 3 -.**
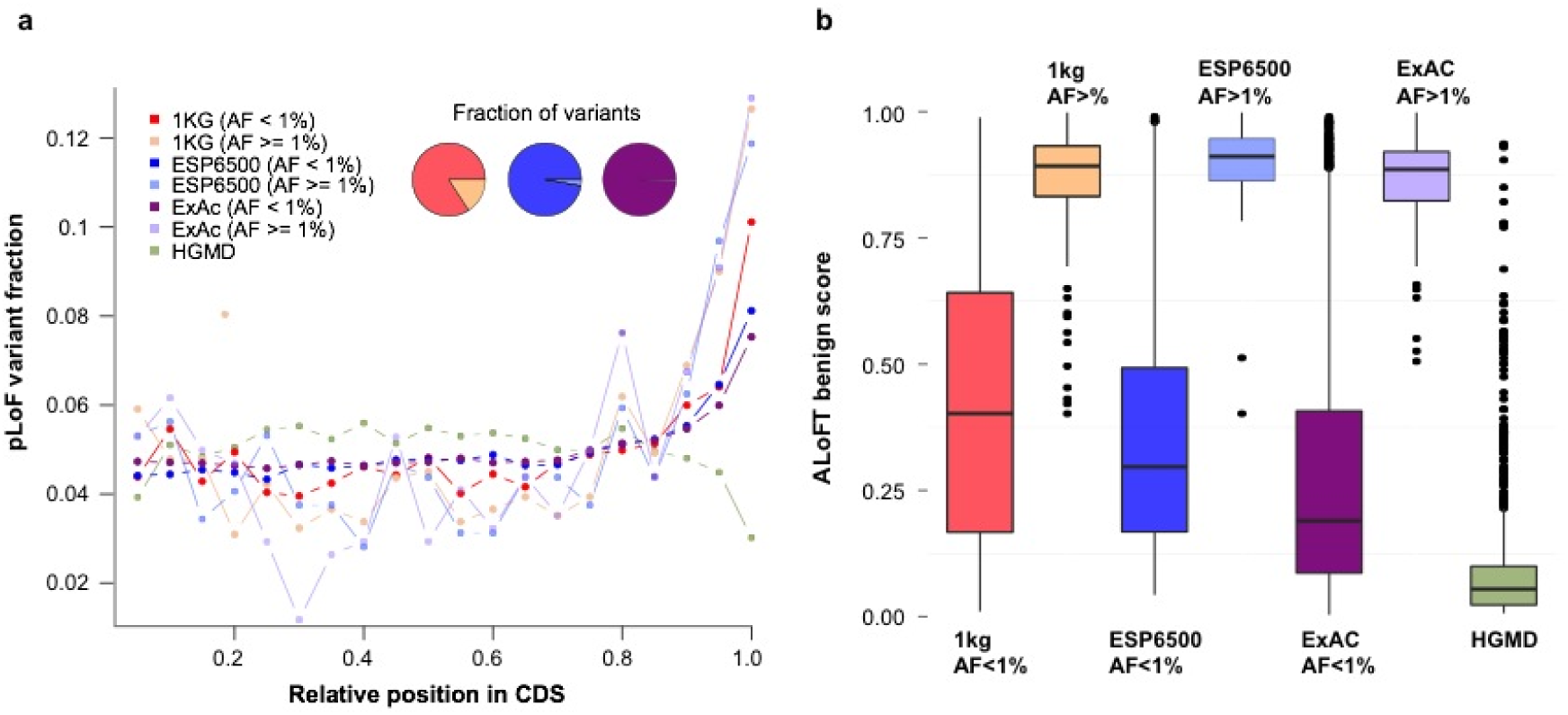
pLoFs in last exons. a) Position of premature stop variants in coding transcripts. Compared to HGMD variants, both common and rare 1KG, ESP6500 and ExAC variants are enriched in the last 5% of the coding sequence. “AF” stands for allele frequency. Variants at allele frequency less than 1% are considered to be rare variants. Variants with at least 1% allele frequency are considered as common. b) Predicted benign scores for premature stop variants in the last coding exons. Training variants are excluded in this plot.

### Application to personal genomes: Understanding pLoFs in an individual genome

The above case studies clearly illustrate the validity of the ALoFT score in elucidating the effect of pLoF variants. In order to estimate the number of pLoF disease alleles in a healthy individual, we applied ALoFT to premature stop variants from the 1KG and ExAC datasets (Supplementary Information). The predicted benign score for pLoFs in these cohorts has a wide range of values (Fig. 4, Supplementary Tables 9,10). Furthermore, due to differences in sequencing coverage and variant calling approaches, the number of potential disease pLoFs per individual varies among datasets. In general, the number increases with higher coverage and larger cohorts where joint variant calling methods result in improved sensitivity in the identification of rare variants. To conservatively estimate a lower bound for per individual statistics (Supplementary Information), we applied a stringent filtering strategy to restrict to high confidence pLoFs. On average, each individual is a carrier of at least two rare heterozygous premature stop alleles that are predicted to be disease-causing in the homozygous state (Supplementary Table 10). Current estimates of the genetic burden of disease alleles (all types of variations, including LoFs) in an individual vary widely, ranging from 1.1 recessive alleles per individual to 31 deleterious alleles^46–50^. In connection with this, it should be noted that the referenced studies are based on diverse methods of identifying variants ranging from targeted panel-based candidate gene studies to whole genome sequencing. The estimation of the number of deleterious pLoF alleles can be affected by a number of confounding factors that include incomplete penetrance of disease alleles, variable expressivity, compensatory mutations, marginal variant calls and imperfect training datasets (Supplementary Information).

**Figure 4 -.**
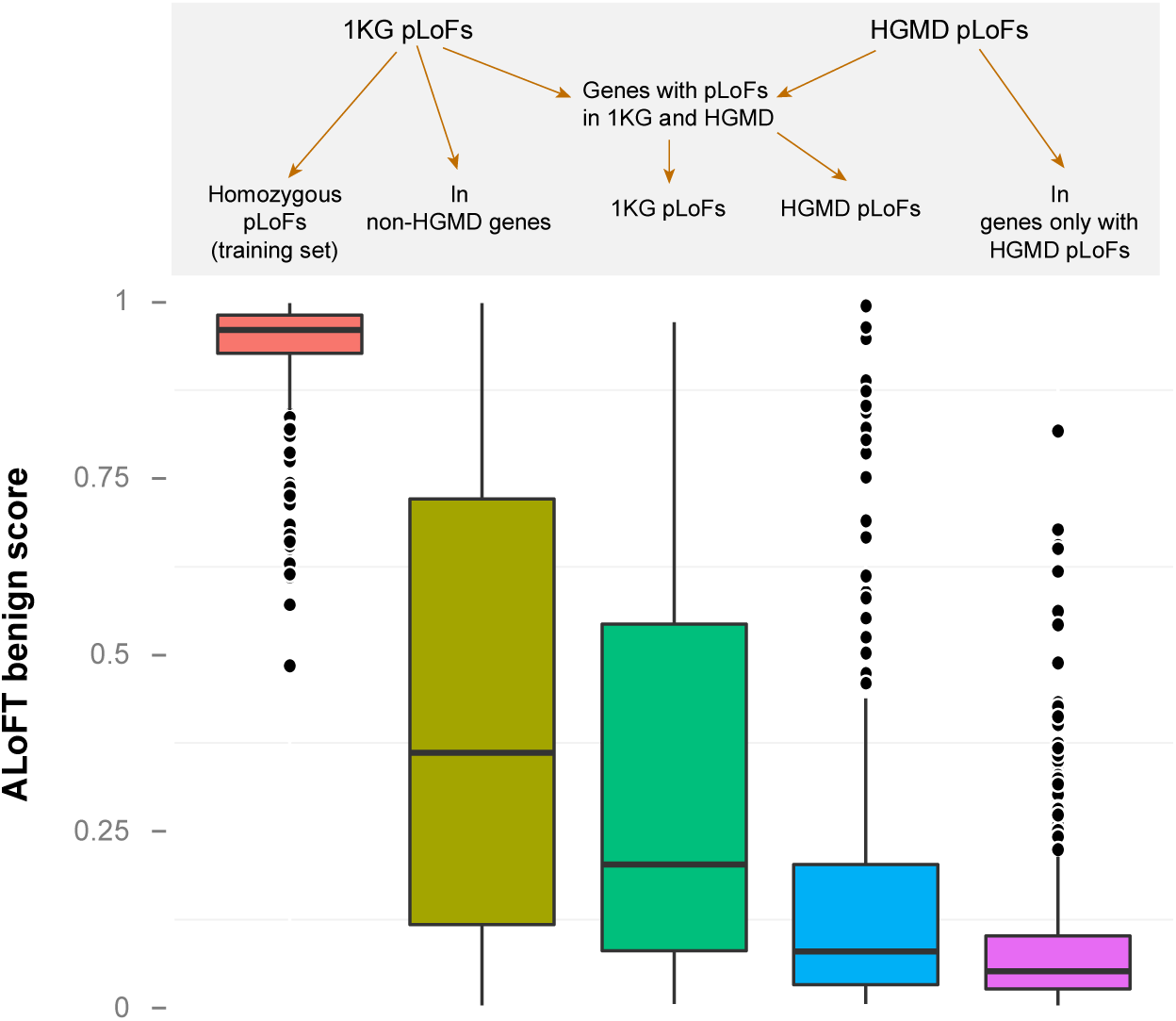
ALoFT classification of 1000 Genomes and HGMD variants. Benign score for premature stop variants in 1KG and HGMD. For this plot, we randomly selected one variant per gene. “Benign pLoFs” set includes homozygous premature stop variants discovered in 1KG. The third (dark green) box plot pertains to premature stop variants in healthy 1KG individuals occuring in disease-causing genes obtained from HGMD. The fourth (blue) box plot pertains to pLoF variants in the subset of HGMD genes where 1KG pLoFs are also seen. “1KG pLoFs in non-HGMD genes” include 1KG variants not in HGMD genes, i.e. non-disease genes. “In genes only with HGMD pLoFs” include HGMD variants in only those disease genes where 1KG pLoFs are not seen.

Next, we looked at premature stop variants in the 1KG cohort in known disease-causing genes. We find that variants in 1KG are more likely to be benign compared to known disease-causing mutations in the same genes (Fig. 4; green vs. blue boxes, p-value: 6.9e-9). Our results provide a possible rationale for this observation. Firstly, variants predicted to be benign in 1KG often affect isoforms that are different from the isoforms containing the disease-causing HGMD variant. This suggests that LoFs in healthy individuals may affect minor isoforms (Supplementary Fig. 9). About 12.4% of premature stop variants in the presumed healthy 1KG individuals in known disease genes and the disease-causing variants in the same genes are on different isoforms. Secondly, some variants predicted to be benign in 1KG occur in the last exon or later in the protein-coding transcript relative to the disease-causing variant in the same transcript. The effect of such variants is possibly the production of truncated proteins that are sufficiently functional. Lastly, a majority of 1KG variants seen in the disease genes are predicted to be disease-causing only if they are homozygous. However, they occur as rare heterozygous variants in the 1KG cohort.

Mutations in HGMD are assumed to be disease-causing. However, some mutations are predicted to be benign by ALoFT (Fig. 4). It is known that disease databases include incorrect disease annotations and common variants and about 27% of variants were excluded by Bell *et al*. in their estimate of carrier burden for severe recessive diseases^46^. Overall, however, only 0.67% of HGMD premature stop mutations are predicted to be benign. Supplementary Fig. 10 shows that most mutations predicted to be benign by ALoFT are seen at higher allele frequencies than those predicted to be in the dominant and recessive classes. Of the 119 pLOF autosomal variants in HGMD predicted to be benign by ALoFT, 32 variants are in Filaggrin, *FLG*. *FLG* LoF mutations are linked to susceptibility to atopic dermatitis, a skin condition leading to eczema^51^. Eczema is a complex trait and the resulting phenotypes are highly variable due to the interplay of environmental and genetic factors^52^. A recent study showed that individuals with bi-alleleic null variants of *FLG* do not always have atopic dermatitis^53^.

A study on British Pakistanis with related parents identified 781 genes containing rare LoF homozygous variants^54^. They found homozygous LoF variants in recessive Mendelian disease genes, however carriers of most of these homozygoys LoF variants do not have the disease phenotype. We applied ALoFT to classify these homozygous LoF variants. Of the 22 variants for which ALoFT provides predictions, 3 are predicted to be benign and none of them were predicted to lead to disease by the dominant mode of inheritance. However, 19 homozygous variants are indeed predicted to lead to disease with a recessive mode of inheritance (Supplementary table 11). The lack of a discernible phenotype could be due to incomplete penetrance of the mutations or due to modifier effects. The penetrance of some disease mutations are also known to be age and sex-dependent^55^. While studies in consanguineous populations have been used to identify recessive disease genes ^56,57^, absence of disease provides an opportunity to look for modifiers in their genetic background.

## Discussion

In summary, we describe ALoFT, a tool for predicting the impact of pLoF variants. In the context of a diploid model, it may be used to determine whether pLoF variants are likely to lead to recessive or dominant disease. Better identification and characterization of pLoF variants has both diagnostic and therapeutic implications. ALoFT allows for the identification and prioritization of high impact putative disease-causing pLoF variants in individual genomes. Integrating benign LoF variants with phenotypic information will help us to identify protective variants which are valuable drug targets^58,59^. Gene functions important for species propagation might actually be deleterious as one ages; thus, LoF variants in such genes provide an intriguing avenue to discover targets for aging-related diseases^60^. Lastly, diseases caused by LoF variants provide opportunities for targeted therapy using drugs that either enable read-through of the premature stop, thus restoring the function of the mutant protein, or NMD inhibitors that prevents degradation of the LoF-containing transcript by NMD^61–67^. This is especially useful in the context of rare diseases where targeting the same molecular phenotype leading to different diseases alleviates the need to design a new drug for each individual disease. Further work will be needed both to correlate the predictions of ALoFT with experimental assays of protein loss of function, and to study the phenotypic impact of heterozygous and homozygous LoF variants in large clinical cohorts.

### Data Availability

The software can be downloaded from aloft.gersteinlab.org. All the ancillary files needed to run the program are included in the download. All the data files discussed in the manuscript have been included as Supplementary data files.

## Acknowledgments

We thank Daniel Spakowicz for comments on the manuscript. This work was supported by grants 5R01GM104371 (US National Institutes of Health/National Institute of General Medical Sciences) to S.B. and D.G.M., and U54HG006504 (Yale Center for Mendelian Genomics) to M.G.

